# MRI R2* captures inflammation in disconnected brain structures after stroke: a translational study

**DOI:** 10.1101/2024.11.18.624105

**Authors:** Ismail Koubiyr, Takayuki Yamamoto, Laurent Petit, Nadège Dubourdieu, Elena Avignone, Elise Cozensa, Chloé Galmiche, Hikaru Fukutomi, Igor Sibon, Vincent Dousset, Michel Thiebaut de Schotten, Aude Panatier, Marion Tible, Thomas Tourdias

## Abstract

Ischemic strokes disrupt brain networks, leading to remote effects in key regions like the thalamus, a critical hub for brain functions. However, non-invasive methods to quantify these remote consequences still need to be explored. This study aimed to demonstrate that MRI-derived R2* changes can capture iron accumulation linked with inflammation secondary to stroke-induced disconnection.

In order to link remote R2* changes to stroke-induced disconnection, we first conducted a secondary analysis of 156 prospectively included stroke patients who underwent MRI at baseline and 1-year follow-up. We mapped fibers disconnected by baseline infarcts to compare the R2* changes over 1 year according to the disconnectivity status in specific thalamic nuclei groups. We also identified the predictors of elevated R2* at 1 year in a multivariate context through linear regressions. In parallel, to understand the biological underpinning of the remote R2* changes, we set up a translational mouse model through photothrombotic induction of focal cortical infarcts or sham procedures in 110 C57BL/6J mice. We explored the mice through combinations of *in vivo* MRI at 72h, 2-, 4-, and 8-weeks, histology, qPCR for gene expression, mass spectrometry for iron concentration quantification, and additional *ex vivo* high-resolution diffusion tensor imaging.

In stroke patients, we found a significant increase of R2* within severely disconnected medial and lateral thalamic nuclei groups from baseline to 1 year. At the same time, no change occurred if these structures were not disconnected. We also showed that the disconnectivity status at baseline was a significant predictor of R2* at follow-up, independently from confounders, establishing a direct and independent relationship between baseline disconnection and the subsequent R2* increase within the associated locations. In mice, we recapitulated the patients’ conditions by observing increased R2* in the stroke groups, specifically within the disconnected thalamic nuclei. Such remote and focal R2* changes peaked at 2 weeks, preceding and correlating with longer-term atrophy at 8 weeks. We established that the remote R2* increase was spatially and temporally correlated with a significant increase of chemically determined iron load bound to ferritin within activated microglial cells.

This study provides critical evidence that R2* is a sensitive marker of inflammation secondary to network disconnection, potentially informing future neuroprotective strategies targeting remote brain regions after stroke.

## Introduction

Ischemic stroke is a major cause of severe long-term disability worldwide.^1^ While revascularization strategies have led to significant improvements in outcomes,^2^ more than one-third of stroke patients who achieve successful recanalization continue to experience persistent difficulties in daily activities and societal roles, despite achieving seemingly excellent functional outcomes (modified Rankin scale 0-1),^3^ even among young adult stroke survivors.^4^ These ongoing challenges are increasingly attributed to post-stroke cognitive and emotional impairments, which affect more than half of stroke survivors.^5,6^ Such impairments in high-order functions may be linked to disrupted interactions between brain regions potentially due to the disconnection of central hubs.^7^

Some critical brain hubs, such as the thalamus or the substantia nigra, are essential to cognitive processes.^8-10^ Although not directly affected by the most frequent middle cerebral artery infarcts as they lie outside this vascular territory, the thalamus and the substantia nigra are highly susceptible to disruption in the extensive thalamo-cortical projections^11^ or nigro-striatal pathway.^12^ This susceptibility means that damage caused by focal infarcts can lead to downstream effects in these hubs, impacting overall recovery. In consequence, alterations in regions remote from the initial infarct may display signs of injury influencing post-stroke outcomes, and present opportunities for neuroprotective interventions.^13^ Quantifying these distant effects *in vivo* becomes critical for comprehensively understanding ischemic consequences and identifying potential biomarkers for future therapeutic trials.

Previously, we^14,15^ and others^16^ have reported secondary increase in R2* values detected by MRI, a quantitative metric sensitive to iron accumulation,^17^ in brain areas remote from the initial infarct. While the anatomical stroke location and remote R2* increase suggest disconnection phenomena,^14,15^ direct evidence linking remote R2* changes to network disconnection is still lacking. Moreover, the biological underpinning of these remote R2* changes after stroke remains poorly understood. The susceptibility effect of iron from hemoglobin and its degradation products following bleeding is the most important contributor to R2* changes in the brain.^18^ Still, R2* might also be sensitive enough to capture high iron content within activated microglial cells in chronic inflammation as shown on MRI-to-pathological correlations from a subset of multiple sclerosis lesions^19^ or in neurodegenerative conditions such as amyotrophic lateral sclerosis,^20^ or progressive multifocal encephalopathy.^21^ Therefore, we hypothesize that similar histopathological changes might underlie *in vivo* observations in stroke patients, reflecting the consequences of network disconnection.

The aim of this study is to demonstrate that remote R2* changes following ischemic stroke reflect an increased iron content associated with inflammation secondary to stroke-inducted disconnection. To address this, we employed a combined approach involving a secondary analysis of clinical data and a translational mouse model.

## Materials and methods

### Clinical data analyses

#### Study population

In order to link remote R2* changes to stroke-induced disconnection, we performed a secondary analysis from a prospective study approved by the institutional ethics committee, with written informed consent from all participants, whose data have already been reported for other analyses (details in supplemental).

The study was conducted in a single center (Bordeaux, France) between June 2012 and February 2015 and enrolled participants according to the following criteria:

Primary inclusion criteria: *(i)* patients older than 18 years old; *(ii)* with a suspected clinical diagnosis of minor to severe supratentorial cerebral infarct [National Institutes of Health Stroke Scale (NIHSS) between 1 and 25]; and *(iii)* confirmed on diffusion-weighted imaging (DWI) at 24–72 h.

Exclusion criteria: *(i)* history of symptomatic cerebral infarct with functional deficit (pre-stroke modified Rankin Scale score ≥1; *(ii)* infratentorial infarct; (*iii)* history of severe cognitive impairment (dementia) or DSMIV axis 1 psychiatric disorders (except major depression); *(iv)* coma; *(v)* pregnant or breastfeeding females; and *(vi)* contraindications to MRI.

Participants underwent 3.0-T MRI scans (Discovery MR 750w; GE Healthcare) at 24-72 hours (baseline) and at 1 year (follow-up) using a consistent protocol including DWI, 3D-T1, and 2D-T2* multi-echo fast gradient echo sequences (details of parameters in supplemental).

We extracted a group of participants from this population without direct thalamic lesion (no thalamic infarct, no thalamic microbleed or hemorrhage) to explore the consequences of thalamic disconnection. We also extracted a group without direct substantia nigra lesion to examine the consequences of disconnection of this other major hub as supplementary analyses (Flowchart in **Supplementary Fig. 1**).

#### Image analyses

Baseline infarcts were delineated on DWI and coregistered to MNI-152 space. We generated disconnection probability maps using BCBtoolkit^22^ which identified likely disconnected tracts by comparing the infarcted regions against tractograms from 163 healthy subjects scanned at 7T, as reported before.^23^ Disconnection maps were thresholded at 0.5, as a standard cut-off,^22^ to include only fibers disconnected in more than half of the healthy subjects used to build the priors.

To assess thalamic disconnection, we overlaid such disconnection maps onto the main groups of thalamic nuclei (lateral, medial, and posterior) from an MNI-152 atlas^24^ and classified patients as having “not disconnected,” “mildly disconnected,” or “severely disconnected” thalamic groups based on the amount of voxel overlap (details in Supplemental and **Supplementary Fig. 2** and 3). We also explored the disconnection of substantia nigra similarly for supplementary analyses.

Then, R2* maps were generated from the voxel-by-voxel mono-exponential fitting of multi-echo T2* images and coregistered to MNI-152 space. We quantified the asymmetry index (AI) of R2* using the non-infarct side as an intrinsic control, as shown below.

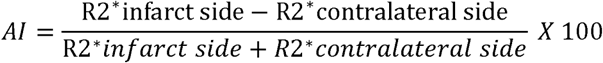

We reported the 95^th^ percentile (AI_95_), a metric reflecting the highest R2* values that has already been used before,^14,15^ and that we compared within each thalamic nuclei group (and substantia nigra) according to their disconnectivity status.

### Animal experiments

#### Animal model

In order to understand the biological substrate of the remote R2* changes, we set up a translational mouse model to recapitulate the observations from the patients.

We used a total of 110 C57BL/6J mice (8 weeks old, 20-25g) from Charles River Laboratories. Experiments were approved by the ethical committee (#28703, #30065 and#13152) and conducted in accordance with the European Directive (2010/63/EU).

We induced a right fronto-parietal cortical infarct sparing the thalamus *via* a photothrombotic ischemia.^25^ We applied a cold light to the surface of the skull, 5 minutes after intraperitoneal injection of a Rose Bengal photosensitive dye (Aldrich Chemical Company, Milwaukee, USA) administered intraperitoneally at a dose of 10 µL/g body weight with a concentration of 5 mg/mL (details in supplemental). Sham-operated mice underwent the same surgical procedure and received the injection of Rose Bengal but without light exposure.

Stroke was confirmed by MRI at 72h. Then, analyses were targeted at 2-, 4-, and 8-weeks post-infarct among three groups (**Supplementary Fig. 4**):

- **Group 1**: Mice underwent sequential *in vivo* MRI before being sacrificed at 2-, 4-, and 8-weeks, followed by paraformaldehyde fixation for histological analyses.
- **Group 2**: Mice were sacrificed at 2-, 4-, and 8-weeks with fresh brain extraction for qPCR and mass spectrometry. All mice in this group also underwent *in vivo* MRI at a single time point before sacrifice, and these additional MRI data were pooled with those from Group 1.
- **Group 3**: Mice were examined with *ex vivo* high-resolution diffusion tensor MRI.

#### In vivo MRI acquisitions and analyses

Mice (n=69 from group 1 and group 2) underwent *in vivo* MRI on a 7T Bruker Biospec system at 72h, 2-, 4-, and 8-weeks, including DWI, 3D-TrueFISP, and 3D-multi echo T2* sequences (parameters in supplemental). Infarcts were outlined on DWI at 72h using 3D-Slicer. As in patients, R2* maps were generated and coregistered to a detailed mouse thalamic atlas^26^ to quantify AI_95_ in specific nuclei and automatically extract their volumes (details in supplemental).

#### Biological analyses

Group 1 of mice (n=27) was sacrificed post-MRI for histological analyses. We stained for microglial cells (Iba1), astrocytes (GFAP), neurons (NeuN), and ferritin light chain. Immunoreactivity was quantified from images collected on a NanoZoomer microscope (40x objective) that captured full brain images through mosaic for direct comparison with MRI. Confocal microscopy (Leica DMI 6000) was also used to assess ferritin colocalization. Details of the procedures are in the supplemental.

Group 2 of mice (n=42) was sacrificed post-MRI for fresh brain extraction, followed by laser-micro-dissection of the thalamus. Briefly, brains were cut using 50 µm thick coronal sections on a freezing microtome (CM3050 S Leica) at -22 °C to prevent RNA degradation. Frozen sections were fixed in a series of precooled ethanol baths and stained with cresyl violet. Subsequently, sections were dehydrated, and a laser capture microdissection was performed using a P.A.L.M MicroBeam microdissection system at 5x magnification. The whole thalamus was captured to maximize the amount of material and because the isolation of specific nuclei was challenging to conduct in such conditions. The material was split into separate adhesive caps to quantify gene expression and iron concentration in the same animal.

For gene expression, RNA was processed and analyzed according to an adaptation of published methods^27^ (details in supplemental) to quantify RNA expression of the complement receptor 3 subunits (CD11b and CD18) that are only expressed by microglia cells,^28^ but also heat shock proteins (HsP1) that are typically upregulated in response to oxidative stress,^29^ and ferritin that is the main iron storage protein (Ftl1).^30^

For chemically determining iron concentration, we used inductively coupled mass spectrometry as a gold standard method that measures the total iron content regardless of its specific chemical form or the protein to which it is bound. We expressed the results in µg of iron per gram of dried brain tissue (details in supplemental).

#### Ex vivo MRI acquisitions and analyses

Group 3 of mice (n=41) underwent *ex vivo* long-scan acquisition (9 hours) on the 7T Bruker for high-resolution diffusion tensor imaging (parameters in supplemental). It included n=20 healthy mice and n=21 stroke mice explored at 2-, 4-, and 8-weeks post-stroke. We computed n=20 whole brain tractograms from the healthy mice which we filtered to extract the streamlines connecting the infarct region to the specific thalamic nuclei in the right hemisphere. After concatenating the 20 thalamo-cortical tracts, we obtained a right thalamo-cortical template which was mirrored across the interhemispheric plane to get a bilateral version. Then, in the n=21 stroke mice, we could measure fractional anisotropy changes along different positions of this template, ipsi- and contralateral to the infarct, at 2-, 4-, and 8-weeks post-stroke. More details are in the supplemental.

### Statistical analyses

In patients, AI_95_ was compared between baseline and 1 year in each group of thalamic nuclei based on their disconnection status (not present, mild, severe) using the Wilcoxon signed-rank test. We also conducted voxel-based comparisons using the Brunner-Munzel test followed by false discovery rate correction. The association between AI_95_ at 1 year and disconnection was also assessed using a linear multivariate model adjusted for age, gender, baseline AI_95_, and infarct volume as potential confounders.

In animals, AI_95_ within specific thalamic nuclei was first compared over time and between groups (stroke *vs.* sham) using 2-way ANOVA followed by post-hoc Tukey tests with p-values adjustment for multiple comparisons. AI_95_ within specific thalamic nuclei and their volumes were also correlated using Spearman’s test. Histological quantifications were compared between stroke and sham groups using repeated Wilcoxon tests and Holm-Šídák correction for multiple comparisons. Fractional anisotropy was compared according to the position along the thalamocortical template and over time using 2-way ANOVA.

Analyses were conducted with GraphPad-Prsim (version 10.2.3) and R (version 4.0.5).

## Results

### Remote R2* increase and disconnection in patients

A total of 156 stroke participants (**Supplementary Fig. 1**) have been prospectively explored at baseline and after 1 year and included in the thalamus analysis (**Table 1**). We mapped each participant’s white matter tracts disconnected by the infarct to assess whether disconnections involved specific thalamic nuclei groups. The anterior thalamic group was too small to run such analysis properly, and we focused on the lateral, medial, and posterior thalamic nuclei groups. Similarly, we also analyzed the disconnection status of the substantia nigra (**Supplementary Fig. 1** and **Supplementary Table 1**). Altogether, we could classify the thalamic nuclei and substantia nigra as “not disconnected”, “mildly disconnected”, or “severely disconnected” (**Supplementary Table 2**) to uniquely test the direct association between such disconnection status and remote R2* changes over 1 year.

**Table 1:** baseline characteristics of the participants included in the thalamus analysis.

| Variables | Participants included in the thalamus analysis<br>(n=156) |
| --- | --- |
| Demographics |  |
| Age (y)* | 65 (56, 77) |
| Sex |  |
| M | 108 (69) |
| F | 48 (31) |
| Hypertension | 101 (65) |
| Diabetes mellitus | 26 (17) |
| Active smoking | 72 (46) |
| Body mass index (Kg/m<sup>2</sup>)* | 26.9 (24.1, 29.4) |
| Baseline visit |  |
| NIHSS score* | 3.0 (2.0, 6.0) |
| Time from onset to baseline MRI (h)* | 46 (33, 59) |
| Recanalization procedure (thrombolysis and/or thrombectomy) | 71 (46) |
| Infarct volume (mL)* | 10 (2, 29) |
| Follow-up visit |  |
| Time from onset to follow-up (y)* | 1.0 (0.98, 1.01) |
| NIHSS score* | 1.0 (0.0, 2.0) |
| Modified Rankin scale (mRS)* | 1.0 (0.0, 2.0) |

Except where indicated, data are numbers of patients with percentages in parentheses. \* Data are median, with IQRs in parentheses.

Participants with severely disconnected lateral and medial thalamus showed significant AI_95_ increases from baseline to 1 year (median, -0.88 [IQR, -5.02-4.33] vs. 7.21 [IQR, 1.30-13.8]; p<0.001 and median, -1.19 [IQR, -3.96-2.00] vs. 4.57 [IQR, -1.35-8.29]; p<0.001 for lateral and medial groups respectively) while, importantly, no significant AI_95_ changes were observed if these structures were not or mildly disconnected (**Fig. 1A**). The voxel-based analysis confirmed that clusters of AI_95_ increases from baseline to 1 year were only found in the groups of severely disconnected lateral and medial thalamus (**Fig. 1B**). Similar AI_95_ increases were observed in participants with severely disconnected substantia nigra but not otherwise (**Supplementary Fig. 5**). We found no significant AI_95_ variation according to the disconnection status within the posterior thalamus, but nevertheless, the voxel-based analysis revealed clusters of voxels with significantly higher values in sub-regions of the posterior thalamus only in severely disconnected participants (**Fig. 1B**).

**Figure 1:**
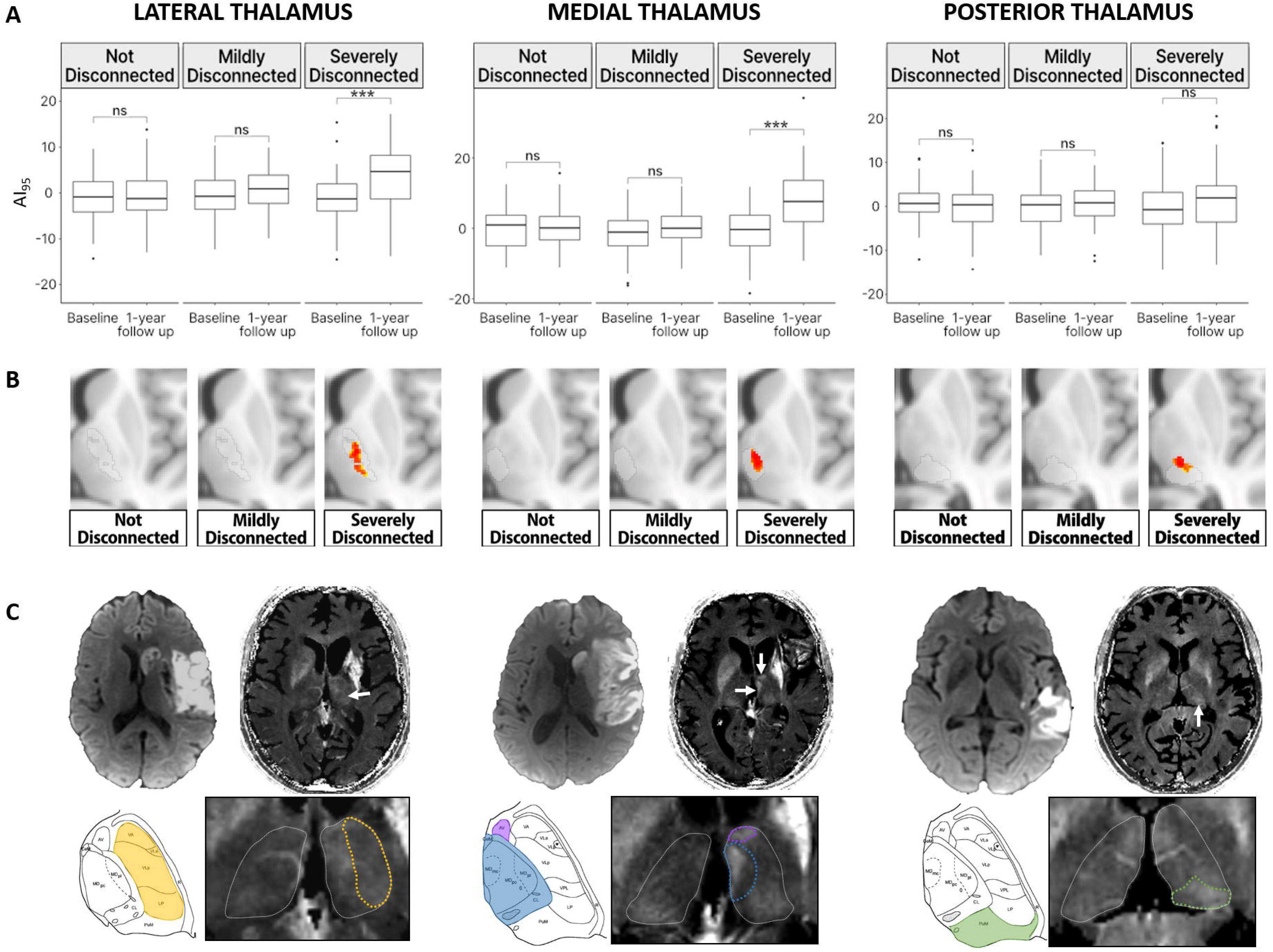
Relationship between disconnectivity and secondary R2* increase at 1-year follow-up in patients. **(A)** shows the evolution of AI_95_ from baseline to 1 year, categorized by disconnectivity status for the lateral, medial, and posterior groups of thalamic nuclei. **(B)** presents the voxel-based comparisons with color-coded clusters that showed a significant increase from baseline to 1 year. **(C)** presents individual examples of severely disconnected cases, showing baseline diffusion images and corresponding R2* map at 1-year follow-up. White arrows point toward focal thalamic R2* increases on the stroke-affected side compared to the contralateral side, magnified below with a delineation of the thalamic groups where the highest asymmetry was measured. For reference, corresponding Morel atlas plates delineate the affected groups of thalamic nuclei. Of note, the focal R2* increase within the medial group (blue) is also associated with a visible increase in the small anterior group (purple).

The qualitative review revealed that focal areas of high R2* could also be identified by simple visual inspection on follow-up MRI, but only for participants with severely disconnected thalamic groups (**Fig. 1C**).

To ensure that the disconnection status was truly a predictor of R2* at 1 year independently from confounders such as baseline AI_95_ or stroke volume, we also ran multivariate linear regression analyses. It showed that disconnection severity was a significant and independent predictor of AI_95_ at 1 year for the lateral (p=0.002) and medial thalamus (p<0.001; **Table 2**), and substantia nigra (p=0.004; **Supplementary Table 3**).

**Table 2:** Multivariable linear regression models for prediction of AI95 at 1 year in each group of thalamic nuclei.

| Variable | B value (95% CI) | P Value |
| --- | --- | --- |
| Disconnection of LATERAL group of thalamic nuclei |  |  |
| Age (+1 y) | -0.04 (-0.09, 0.01) | 0.10 |
| Sex (F) | -0.48 (-1.9, 0.96) | 0.50 |
| AI<sub>95</sub> at baseline (+ 1) | **0.28 (0.16, 0.40)** | **<0.001\*\*\*** |
| Infarct volume (+ 1 mL) | 0.01 (-0.01, 0.02) | 0.30 |
| Disconnection |  |  |
| Mildly disconnected | -0.62 (-2.4, 1.2) | 0.50 |
| Severely disconnected | **2.9 (1.1, 4.7)** | **0.002\*\*** |
| Disconnection of MEDIAL group of thalamic nuclei |  |  |
| Age (+1 y) | **-0.07 (-0.13, -0.01)** | **0.013\*** |
| Sex (F) | -1.0 (-2.6, 0.57) | 0.20 |
| AI<sub>95</sub> at baseline (+ 1) | **0.36 (0.24, 0.48)** | **<0.001\*\*\*** |
| Infarct volume (+ 1 mL) | **0.05 (0.04, 0.07)** | **<0.001\*\*\*** |
| Disconnection |  |  |
| Mildly disconnected | 0.87 (-3.1, 1.3) | 0.40 |
| Severely disconnected | **5.2 (2.9, 7.5)** | **<0.001\*\*\*** |
| Disconnection of POSTERIOR group of thalamic nuclei |  |  |
| Age (+1 y) | 0.01 (-0.04, 0.06) | 0.60 |
| Sex (F) | -1.3 (-2.7, 0.16) | 0.081 |
| AI<sub>95</sub> at baseline (+ 1) | **0.14 (0.01, 0.27)** | **0.038\*** |
| Infarct volume (+ 1 mL) | **0.03 (0.01, 0.04)** | **0.001\*\*** |
| Disconnection |  |  |
| Mildly disconnected | 0.07 (-1.8, 1.9) | 0.90 |
| Severely disconnected | 0.88 (-1.0, 2.8) | 0.40 |

Overall, these results established a direct and independent relationship between baseline disconnection and subsequent R2* increases, specifically in the disconnected structures.

### Dynamic of remote R2* increase in the animal model

As pathological tissues are not collected in stroke patients, we set up a mouse model recapitulating the human observations, which we could use to assess the biological underpinning. The adapted photothrombosis model produced reproducible lesions with restricted diffusion in the sensorimotor cortex (**Supplementary Fig. 6**), leading to high T2 signal at 72 hours, followed by progressive cortical atrophy, resembling typical human cortical stroke (**Fig. 2A**). Infarcts always spared the thalamus. There were no lesions in sham-operated mice.

**Figure 2:**
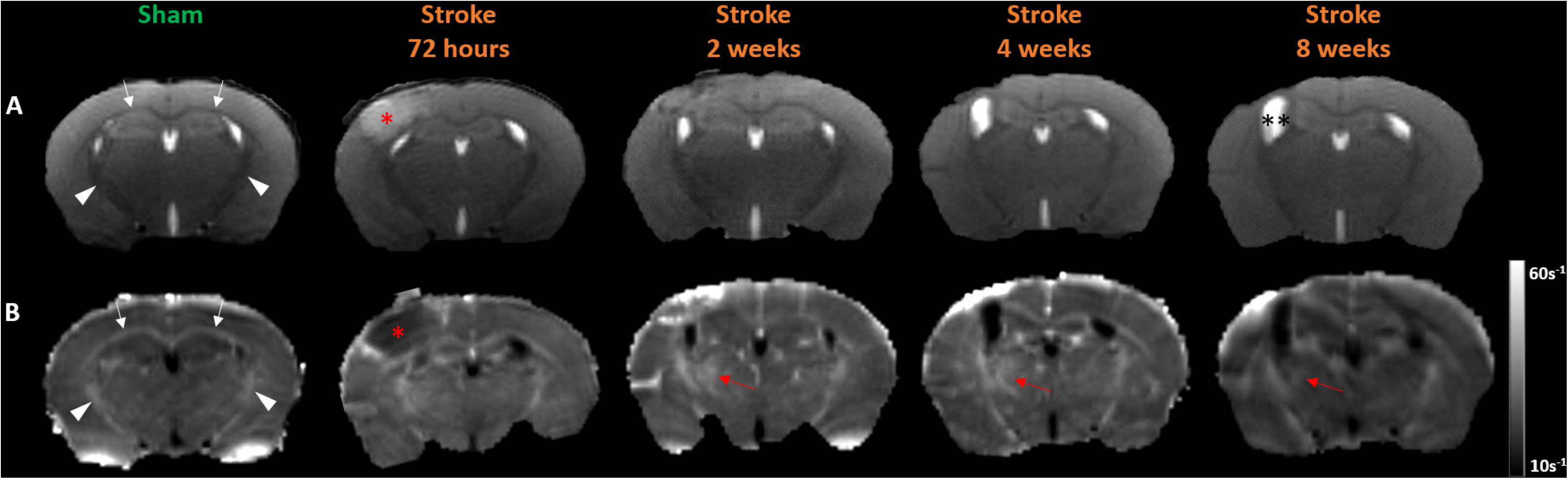
Dynamic evolution of stroke and R2* in the mouse model. **(A)** displays T2-weighted images and **(B)** R2* maps in a sham-operated and a stroke mouse followed longitudinally with repeated MRI at 72h, 2-, 4-, and 8-weeks. The focal infarct, indicated by the red asterisk, is clearly visible at 72 hours and evolves into focal atrophy with passive dilatation of the ipsilateral ventricle (**). R2* maps show higher values within iron-rich myelinated white matter tracts, such as the corpus callosum (white arrows) and internal capsules (white arrowheads). At 2 and 4 weeks, a focal area with R2* increase appears in the thalamus ipsilateral to the stroke and becomes subtler at the 8-week follow-up (red arrows).

Visual inspection consistently revealed focal thalamic R2* increases at 2- and 4-week post-stroke, with subtler changes at 8-weeks (**Fig. 2B**). Co-registration with a thalamic atlas (26) localized R2* increases specifically within the ventral posterolateral (VPL) and ventral posteromedial (VPM) nuclei (**Fig. 3A**). Quantification of AI_95_ within these nuclei showed a significant increase in stroke mice compared to sham (group effect, p=0.019 and 0.009 in VPL and VPM, respectively), with distinct temporal dynamics (group x time interactions p<0.05) peaking at 2 and 4 weeks for the stroke mice (**Fig. 3B**). Additionally, we measured significant ipsilateral volume loss in VPL and VPM in stroke mice compared to sham (group effect, p=0.008 and 0.04 in VPL and VPM respectively), worsening progressively in stroke but not in sham mice (group x time interactions p<0.05) and becoming prominent at 8-week follow-up (**Fig. 3C**). As an additional control, we also identified three distant thalamic nuclei (AND, IDN, LH) with volumes close to those of VPL and VPM but distinct thalamo-cortical projections, and found no significant AI_95_ or volume changes over time in these nuclei in stroke mice compared to sham.

**Figure 3:**
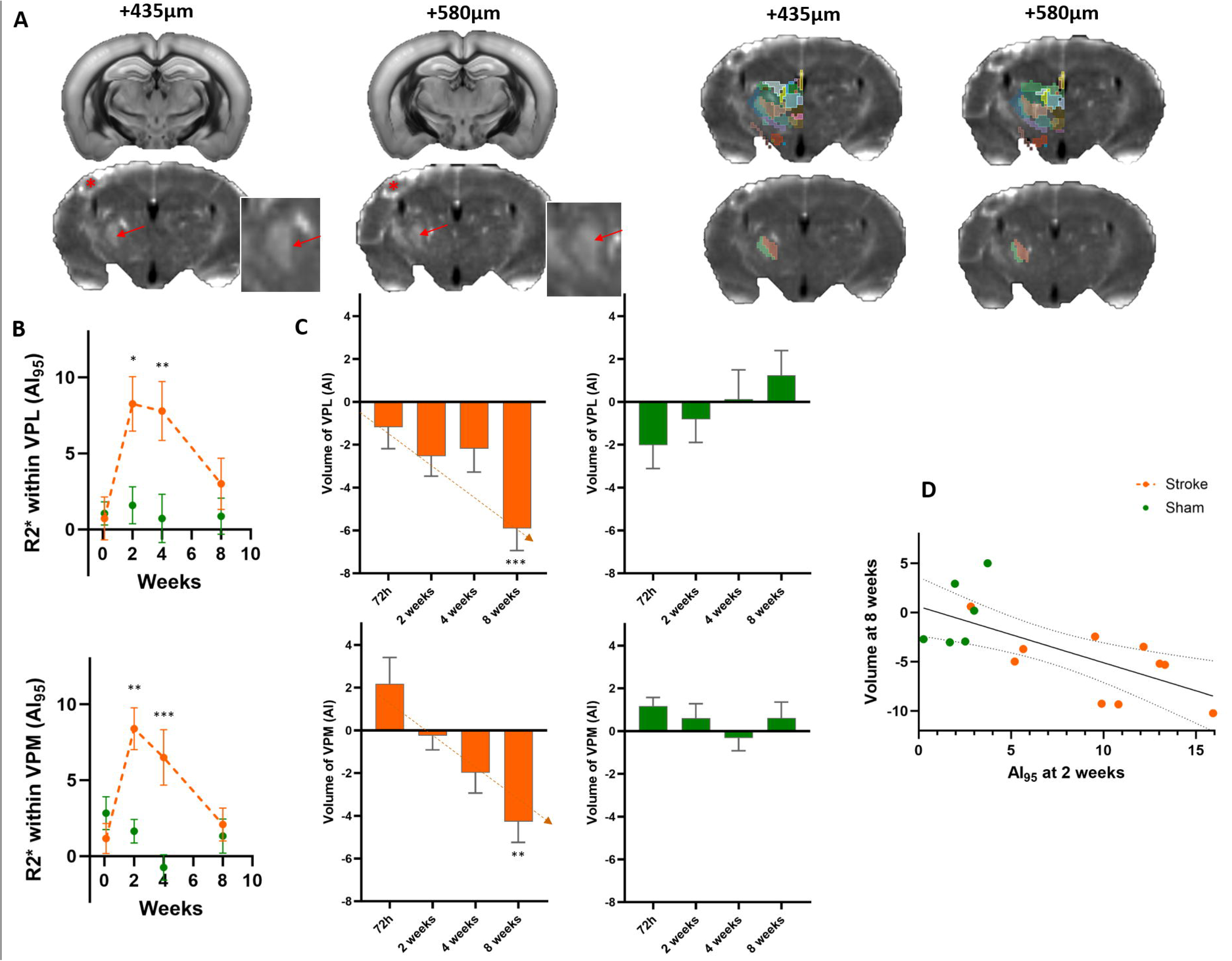
Quantification of R2* and thalamic volumes over time in the mouse model. **(A)** provides an example of R2* map co-registration with the thalamic atlas (26). Two different z-positions are displayed from a stroke-induced mouse at 2 weeks. The stroke is marked with a red asterisk. Focal R2*increases are visually identified (red arrows, magnified in insets). Among all the thalamic nuclei (top row), we observed that the focal R2* increases were precisely located within VPL (green) and VPM (red) nuclei, which are the only two nuclei maintained in the bottom row. **(B)** shows the time course of R2* quantification using AI_95_ and **(C)** presents the time course of VPL and VPM volumes for stroke (orange) and sham-operated (green) mice. In stroke mice, R2*peaks at 2 weeks, while volumes progressively decrease until 8 weeks. **(D)** shows a significant inverse correlation between R2* at 2 weeks within VPL and VPM and the volumes of these nuclei at 8-week follow-up. Error bars indicate mean ± SEM. *, p≤0.05; **, p≤0.01; and ***, p≤0.001 from Tukey post-hoc tests of stroke vs. sham with p values adjusted for multiple comparisons.

The mice followed longitudinally with repeated MRIs until 8 weeks provided the unique opportunity to directly correlate R2* and volume changes over time within the same animals. Such analysis revealed that the R2* increases in VPL and VPM at 2 weeks (the peak of the time-course curve) were significantly correlated with subsequent atrophy measured at 8 weeks (**Fig. 3D**; r, -0.72; [95%CI, -0.89, -0.33]; p=0.0024).

Overall, these results revealed that remote and focal R2* increase can be modeled through experimental cortical stroke in mice, resembling closely the human pattern, and they showed that R2* increase preceded and predicted longer term atrophy.

### Biological substrate of remote R2* increase

We ran histology and molecular biology to elucidate the modifications associated with the remote R2* increase. Histological analyses at 2-, 4-, and 8-weeks post-stroke, conducted immediately after MRI scans, showed glial cell activation in regions delineating precisely VPL and VPM in close correspondence with the observed R2* increases in MRI (**Fig. 4A**). We also found astrocytic reactivity with increased GFAP staining at all time points, alongside the microglial activation evidenced by larger cell bodies and thicker processes that translated into significant Iba1 staining increases (**Fig. 4B and 5A**). Notably, the mRNA expression level of the microglia-specific receptor of the complement (CD11b and CD18 sub-units), which is a marker of microglial activation, peaked at 2- and 4-weeks, which also came with a significant mRNA increase of heat shock protein-1 at 2 weeks (**Fig. 5B**). Neuronal count decreased progressively until marked reduction, aligning with MRI-measured atrophy at 8 weeks (**Fig. 5A**).

**Figure 4:**
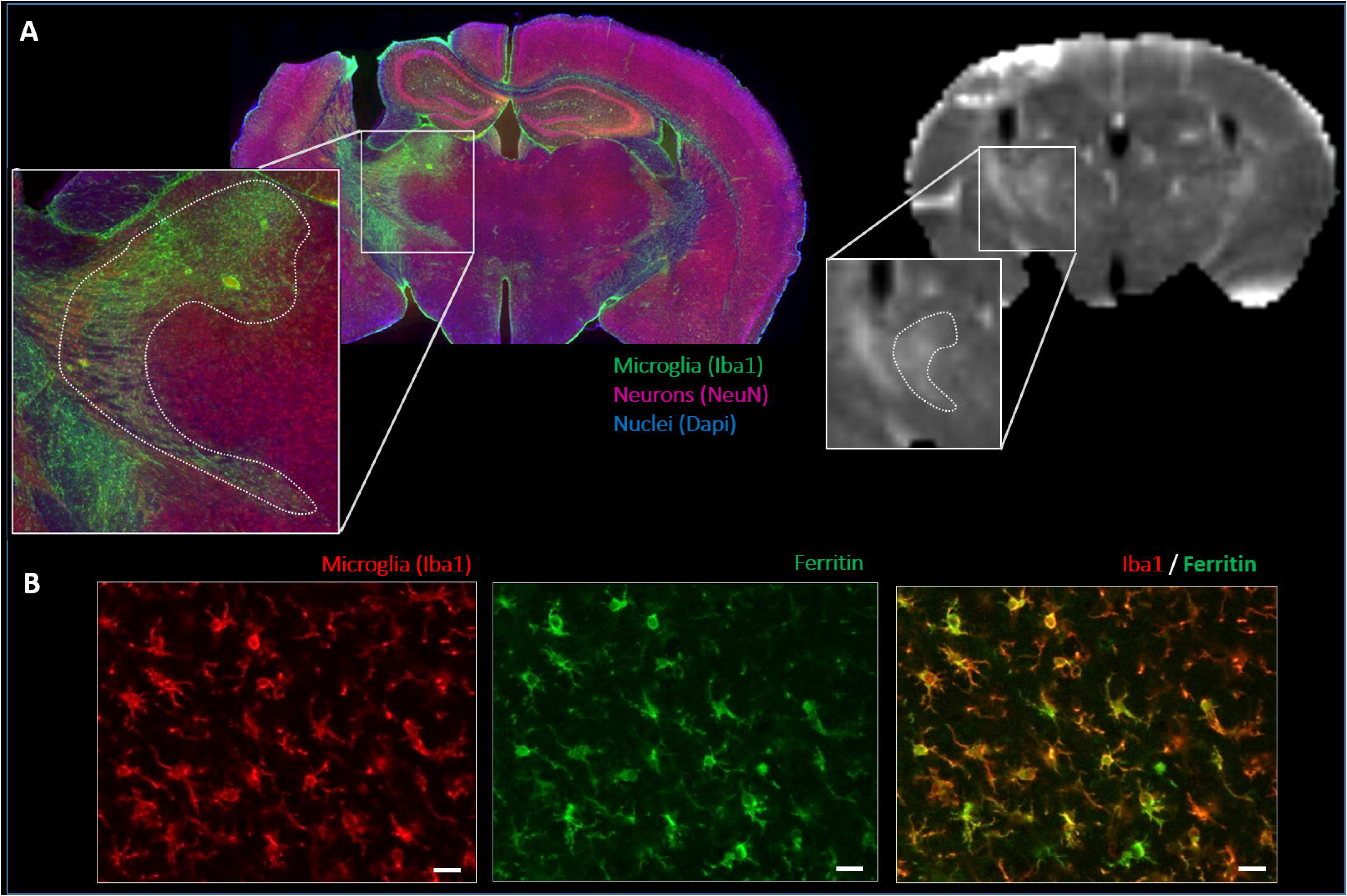
Histological-to-MRI comparison. **(A)** shows immunostaining of microglial cells (green) and neurons (red) with blue nuclear counterstain (DAPI) alongside the corresponding R2* map acquired prior to sacrifice. Enlarged views highlight microglial activation within the ventral posterolateral (VPL) and ventral posteromedial (VPM) thalamic nuclei (dotted lines), spatially correlated with the focal R2*increase. **(B)** displays higher magnification confocal images of microglial cells (red) and ferritin light chain (green), demonstrating ferritin localization within activated microglia, which exhibit enlarged cell bodies and thicker processes. The scale bar is 100 µm.

**Figure 5:**
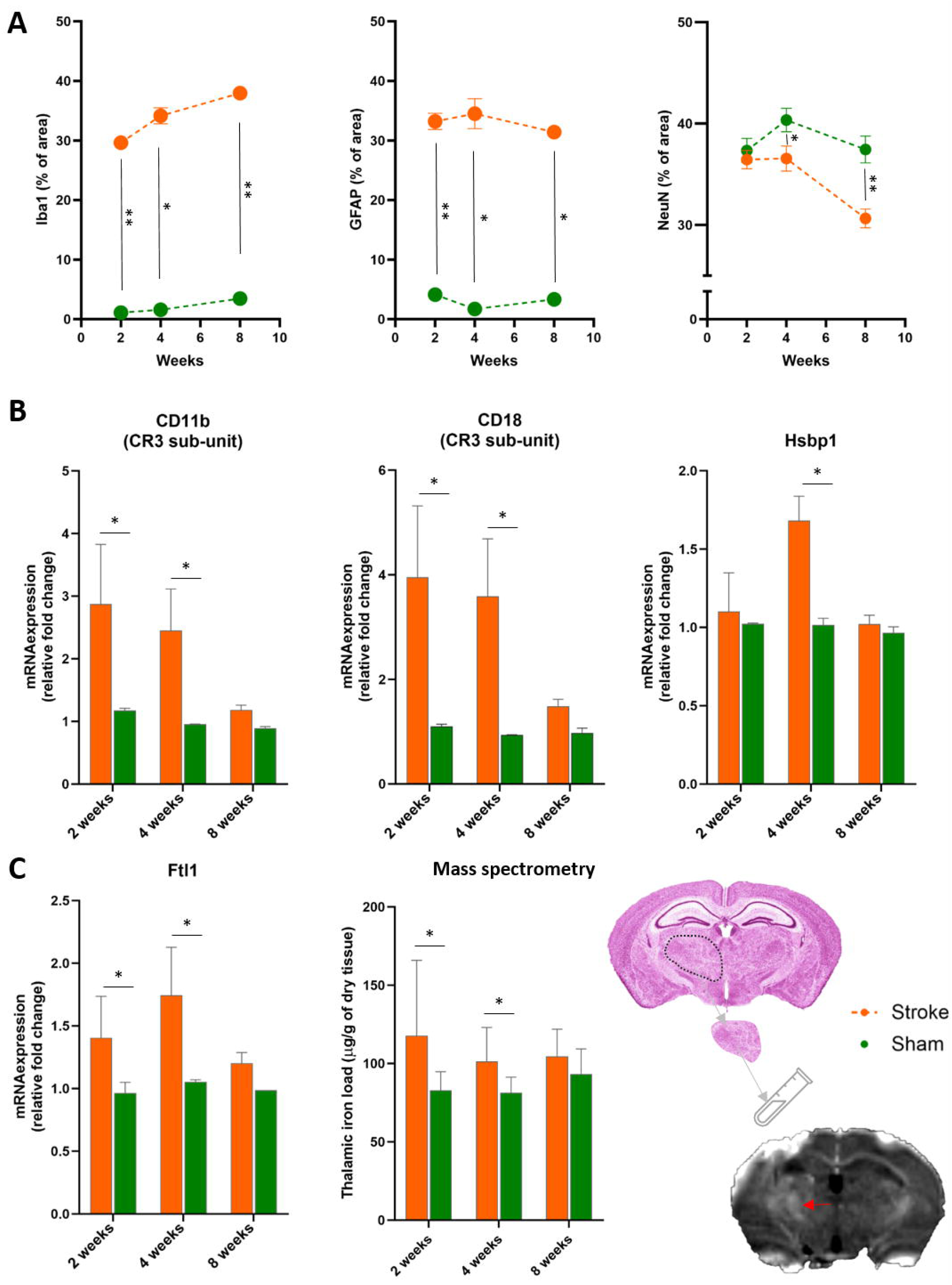
Quantification of immunostaining, gene expression, and iron concentration. **(A)** presents the semi-quantitative analysis of microglia (Iba1), astrocytes (GFAP), and neurons (NeuN) immunoreactivity over time, expressed as the percentage of area stained above an identical threshold on the stroke side compared to the contralateral side. **(B)** shows relative gene expression levels in stroke (orange) compared to sham (green). **(C)** displays iron-related markers with ferritin gene expression and iron concentration from mass spectrometry, both extracted from laser-capture microdissections of the thalamus where the high R2* spots (red arrow) have been observed. Error bars indicate mean ± SEM. *, p≤0.05; and **, p≤0.01 from Wilcoxon tests and Holm-Šídák correction for multiple comparisons.

The specific quantification of iron via mass spectrometry in laser microdissected thalamus tissue ipsilateral to stroke showed significantly increased total iron load at 2 and 4 weeks that didn’t reach significance at 8 weeks (**Fig. 5C**). The mRNA expression of ferritin, the primary iron storage protein, was significantly increased at these time points (**Fig. 5C**), with confocal microscopy showing ferritin localization within the activated microglial cells (**Fig. 4B**).

In total, these results established that the remote R2* measured *in vivo* was spatially and temporally correlated with higher iron load mainly bound to ferritin within pro-inflammatory microgial cells.

### Remote R2*increase and disconnection in mice

We collected high-resolution diffusion MRI to ensure that the remote VPL and VPM modifications described above during the follow-up were directly related to thalamo-cortical projection disruption. In healthy mice, we extracted a white matter pathway connecting the cortical stroke site to ipsilateral VPL and VPM within the thalamus, indicating a direct connection between these regions. This tract passed below the lateral ventricle, crossed the distal part of the corpus callosum, ran between the caudate and putamen before reaching specifically the VPL and VPM thalamic nuclei (**Fig. 6A**) corresponding well with specific dye injection within VPL and VPM from the Allen Brain Atlas (**Supplementary Fig. 7**).

**Figure 6:**
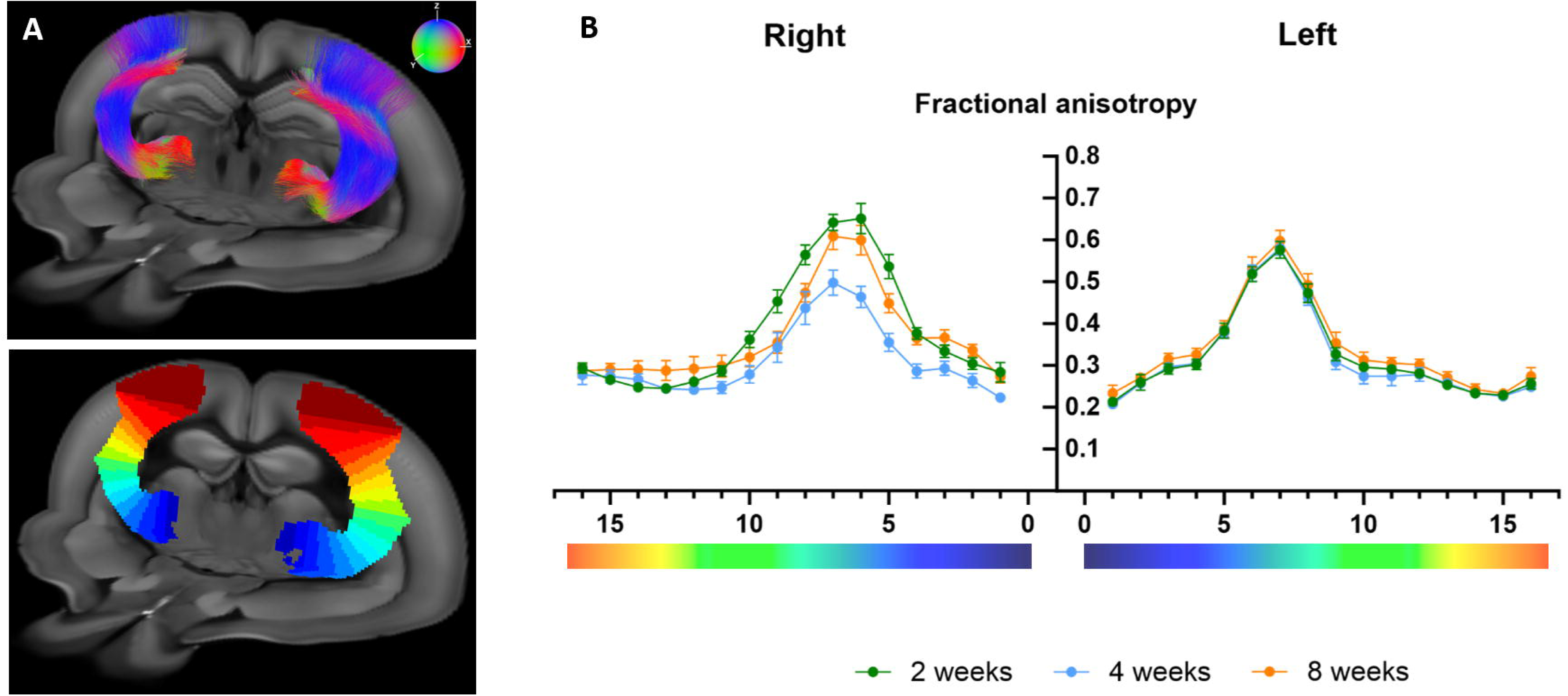
Thalamo cortical fibers connecting stroke site to VPL and VPM from ex vivo diffusion MRI. **(B)** shows a 3D projection of MR tractography between the stroke site and VPL / VPM. Colors indicate tract directions: green for rostro-caudal, red for left-right, and blue for dorsal-ventral. The bottom panel shows segmentations of these thalamocortical tracts in 16 evenly distributed segments color-coded by position. **(B)** depicts fractional anisotropy quantification in stroke mice on the stroke (right) and contralateral side (left) at 2, 4 and 8 weeks, with quantification conducted within the 16 segments indicated on the abscissa and color-coded by position as in A.

We used this pathway as a template to quantify microstructural changes in stroke mice. On the contralateral side (left), fractional anisotropy varied significantly according to the position along the tract (position effect, p<0.0001) in line with the anatomical characteristics of this bundle, but showed no time-dependent changes following stroke. On the stroke side, we found significant time-dependent variations (p=0.003) that differed according to the position within the tract (time x position interaction, p<0.001). Notably, at the higher anisotropy position close to the tract midpoint, fractional anisotropy decreased significantly from 2 to 4 weeks which was followed by partial recovery at 8 weeks, though not reaching 2-week levels (**Fig. 6B**).

These results confirmed that the R2* changes within VPL and VPM were related specifically to stroke-induced disconnection as the microstructure of the direct connecting pathway was significantly altered.

## Discussion

Brain functions emerge from dynamic interactions between areas connected by white matter tracts.^7^ Ischemic strokes disrupt these connections, leading to remote effects that are challenging to quantify *in vivo*. In this translational study, we demonstrate that remote R2* increases occur specifically in disconnected structures, such as thalamic nuclei or substantia nigra. We also showed that these R2* increases reflect iron-rich microglial activation, which precedes and predicts secondary atrophy. This suggests that iron imaging can provide a comprehensive view of remote stroke consequences and potentially inform future neuroprotective strategies.

Remote changes in regions distant from focal cortical strokes, particularly in the thalamus or the substantia nigra, have been reported for many years through a variety of imaging approaches^31-34^ with R2* having the potential to capture the long-term modifications.^14,15^ Traditionally attributed to anterograde or retrograde degeneration based on anatomical presumptions, these changes have lacked direct evidence linking them to fiber disconnection. By mapping focal infarct lesions onto high-resolution connectomes,^22,23^ we could quantitatively measure R2* changes for the first time in specific hub regions that have been disconnected or not according to the initial infarct location. This demonstrated that the delayed R2* reaches significant increase only following substantial disconnection. The posterior thalamus behaved differently from the lateral and medial group or the substantia nigra which will require further explorations. One potential explanation could be that the type of secondary degeneration depends on the amount of afferent versus efferent pathways. Interestingly, the pulvinar shows mainly dense efferent projections (output) to cortical regions while the medial and lateral thalamic nuclei groups, as well as the substantia nigra, have a more balanced afferent (input) and efferent (output) connectivity pattern^8,12^ that could produce more extensive degeneration.

Additionally, using diffusion MRI in mice, we reconstructed a thalamo-cortical pathway between experimentally induced cortical infarcts and altered thalamic nuclei, directly demonstrating microstructural alterations within bundles connecting affected nuclei. The reduction in fractional anisotropy at 4 weeks aligns with axonal fragmentation, as seen in spinal cord degeneration^35^ while subsequent glial proliferation and cellular debris may contribute to the partial recovery of fractional anisotropy observed at 8 weeks.^36^

The translational approach in mice further evidenced that R2* captures inflammatory responses in disconnected regions. Histological analyses^37^ and *in vivo* PET imaging studies with tracers for the peripheral benzodiazepine receptors^38^ have shown that microglial activation is a key feature of secondary injuries. We confirmed strong microglial activation markers in the disconnected VPL and VPM nuclei, which correlated tightly with increased R2* on MRI, in both spatial and temporal dimensions. Mass spectrometry data indicated that elevated R2* values reflect increased total iron content, likely bound to upregulated ferritin within activated microglial cells. This finding aligns with paramagnetic effects seen in amyotrophic lateral sclerosis^20^ and progressive multifocal leukoencephalopathy,^21^ where iron-rich microglial cells have been implicated in specimen analyses. Our observation is also reminiscent of a particular histological subtype of multiple sclerosis lesions called the “chronic active lesions” that are characterized by a demyelinating core surrounded by a dense rim of activated microglial cells.^39^ Interestingly, the so called “paramagnetic rim lesion, PRL” on susceptibility sequences emerged as an appealing *in vivo* MRI marker for these chronic active lesions^40^ with the source of the signal of the rim being molecular iron sequestered intracellularly and bound to ferritin within pro-inflammatory macrophages and microglia.^19^ Here, we extend the role of R2* as a marker of neuroinflammation in secondary degeneration post-stroke. Imaging allowed us to trace the inflammatory response over time non-invasively showing, in mice, a delayed onset, peaking between 2-4 weeks and declining at 8 weeks, consistent with more complex PET-based or invasive histological analyses in other models.^41,42^ Importantly, R2* was sensitive enough to capture subtle inter-individual differences despite the standardized induction of the model, and served as an earlier biomarker than atrophy, representing the late and irreversible stage of neurodegeneration. The correlation between R2* and subsequent atrophy implies a potential causal relationship between activated microglia and neurodegeneration, potentially involving oxidative stress mechanisms, supported by the upregulation of heat shock proteins. This is further corroborated by the recent identification of a neurodegenerative subtype of microglia (CD11c^+^) in the disconnected thalamus.^43^ Our findings in stroke patients closely parallel those in the mouse model, indicating that R2* iron imaging could serve as a valuable biomarker to quantify and monitor remote stroke consequences, preceding neuronal loss as measured from flumazenil PET^34^ or atrophy.^44^

In spite of these promising results, our study was limited by the absence of behavioral data, which made it underpowered to demonstrate behavioral correlates to remote changes. However, previous works have linked delayed injury to cognitive and emotional deficits.^14,45^ In patients, we relied on indirect disconnectome methods rather than direct diffusion tensor imaging, which may have limited our ability to perfectly assess individual disconnection. However, such indirect disconnectivity method has been extensively validated.^22,23,46^ In animals, we haven’t identified iron directly in histological sections through Perls Prussian blue staining, but we used the gold standard mass spectrometry in microdissected tissue that is the only able to provide true quantitative measurement. Finally, the metabolic pathways of iron sequestration haven’t been explored and whether iron accumulation is causally involved in the associated neurodegeneration, for instance through ferroptosis,^47^ is an ongoing area of investigation which could be the focus of future research.

In conclusion, a remote increase in R2* in disconnected brain structures following stroke represents neuroinflammation that precedes neuronal loss. This imaging biomarker could be crucial for advancing neuroprotective trials aimed at preserving brain networks.^48^

## Data availability

The tabulated data that support the findings of this study are available from the corresponding author, upon reasonable request from a qualified investigator.

## Acknowledgements

We thank the contribution of the AEM2 Platform (Martine Ropert-Bouchet, University of Rennes 1 / Biochemistry Laboratory, University Rennes Hospital) for mass spectrometry and the contribution of the PUMA platform (Thierry Lesté-Lasserre and Marlène Maître, Neurocentre Magendie, INSERM U1215) for laser microdissection and quantitative PCR. The MRI acquisitions were performed at the Institute of Bioimaging (UAR 3767), and the microscopy was done in the Bordeaux Imaging Center, a service unit of the CNRS-INSERM and Bordeaux University, both members of the national infrastructure France BioImaging supported by the French National Research Agency (ANR-10-INBS-04).

