## supplemental for "MRI R2* captures inflammation in disconnected brain structures after stroke: a translational study"

**Supplemental materials and methods**

**Study population**

Participants were prospectively recruited as part of a cohort study called BBS for “Brain Before Stroke”. The primary objective of this cohort was to investigate the impact of the modifications outside the infarct itself on stroke outcomes, such as those related to small vessel disease. Analysis of this objective has been reported by Coutureau *et al*. (1). We have already used this cohort to demonstrate the presence of increase R2* outside the infarct region (2, 3). The present paper is a follow-up analysis aimed at examining the relationship between disconnection patterns and delayed R2* increases, which has not previously been explored. Additionally, this cohort has been utilized in prior lesion-symptom mapping studies (4, 5), though those analyses are unrelated to the present work.

The imaging protocol included diffusion weighted images (DWI), 3D-T1-weighted images, and 2D-T2* multi echo fast gradient echo images with the following parameters:

- For DWI: 38 slices; repetition time, 9000 ms; echo time, 76 ms; slice thickness, 4 mm; matrix, 128 x 128; field of view, 240mm x 240 mm; b values, 0 and 1000 s/mm^2^.
- For the 3D T1 inversion-recovery-prepared fast spoiled gradient echo sequence: 196 sagittal slices; repetition time, 8.60 ms; echo time, 3.27 ms; inversion time, 450 ms; flip angle, 12°; slice thickness, 1 mm; matrix, 256 x 256; field of view, 240mm x 240 mm.
- For the 2D multiecho fast gradient-echo sequence: 28 slices; repetition time, 775 ms; eight echo times at 4.3, 8.7, 13.0, 17.4, 21.7, 26.1, 30.5 and 34.8 ms; flip angle, 20°; slice thickness, 4 mm; matrix, 320 x 320; field of view, 240mm x 240 mm.

Out of 428 recruited patients, 156 were included for analyses focusing on the thalamus, and 177 were included for analyses focusing on the substantia nigra, according to the flowchart in **Supp-Figure-1**.

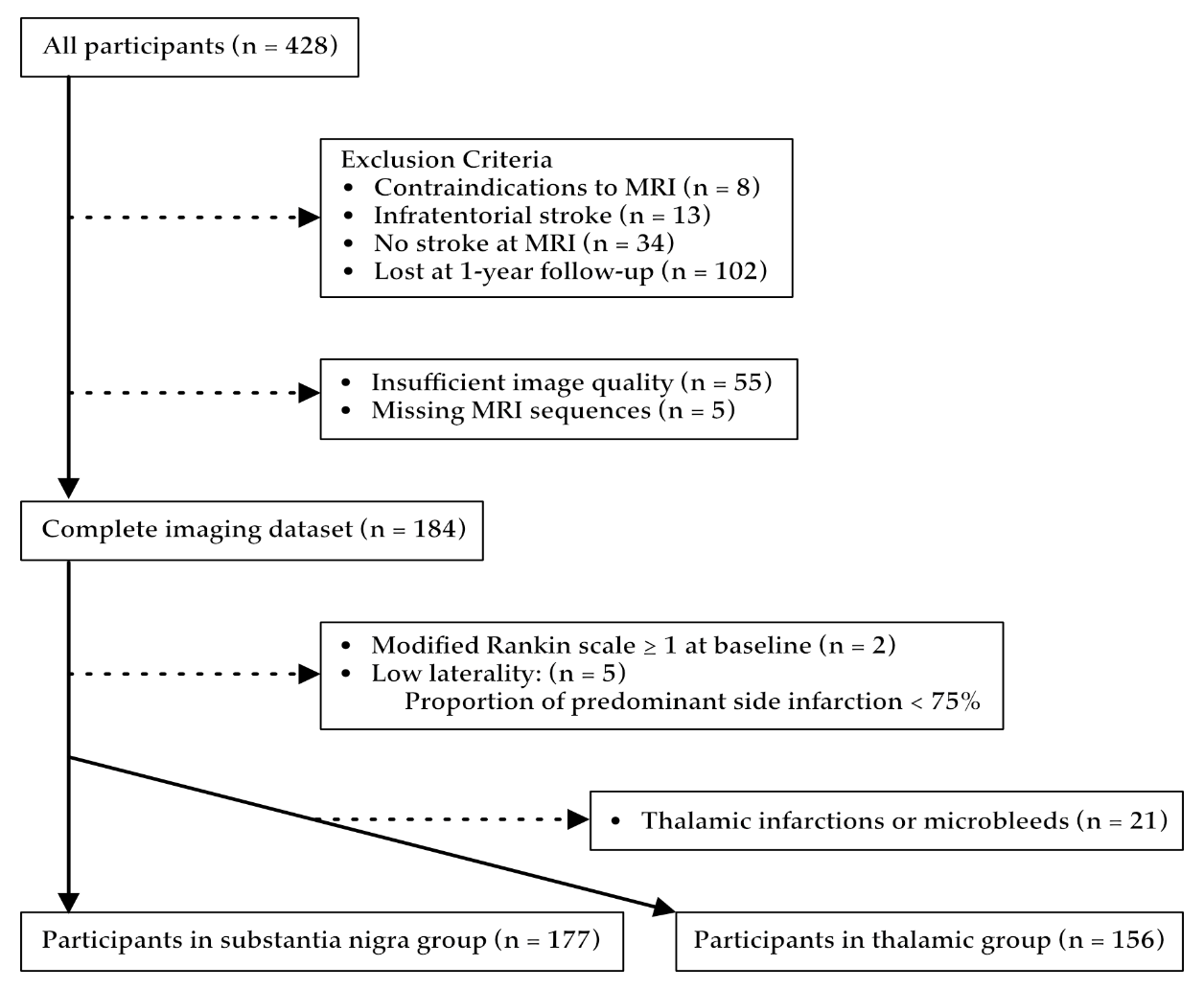

**Supp-Figure 1:** Flowchart of patient inclusion and exclusion

**Image analyses in patients**

To define the disconnection status of remote areas, we assessed the overlap between binarized disconnectivity maps and masks of thalamic nuclei groups (6) (**Supp-Figure-2, upper panel**) and substantia nigra (7) (**Supp-Figure-2, lower panel**), all available in MNI-152 space.

The disconnectivity maps represent the fibers likely to be disconnected in a given stroke patient, based on the premise that more than half of the 163 healthy participants from the Human Connectome Project (8) would exhibit fiber disconnection by an infarct in the same location (9). Therefore, the maps provide a voxel-by-voxel probability of disconnection, ranging from 0.5 to 1, excluding values below 0.5.

For thalamic nuclei, we utilized an atlas that we constructed from 7 Tesla MR images (6) taking advantage of an optimized sequence, called white matter nulled MPRAGE and providing highly contrasted images, to manually delineate thalamic nuclei (10). We combined the thalamic nuclei into 4 main groups following the definitions from Morel atlas (11) as follow:

- Anterior group: anteroventral (AV)
- Lateral group: ventral posterolateral (VPL), ventral lateral anterior (VLa), ventral lateral posterior (VLp), and ventral anterior (VA)
- Medial group: mediodorsal (MD), centromedian (CM), and habenula (Hb) (part of the epithalamus but comprised in the paraventricular complex and located at the infero-postero-medial part of the medial group and so very close to the 3rd ventricle)
- Posterior group: pulvinar (Pul), medial geniculate nucleus (MGN), and lateral geniculate nucleus (LGN)

For the substantia nigra, we used a probabilistic atlas coregistered to MNI-152 and freely available (7).

**Supp-Figure 2:** Schematic shows the pipeline to estimate the disconnectivity status
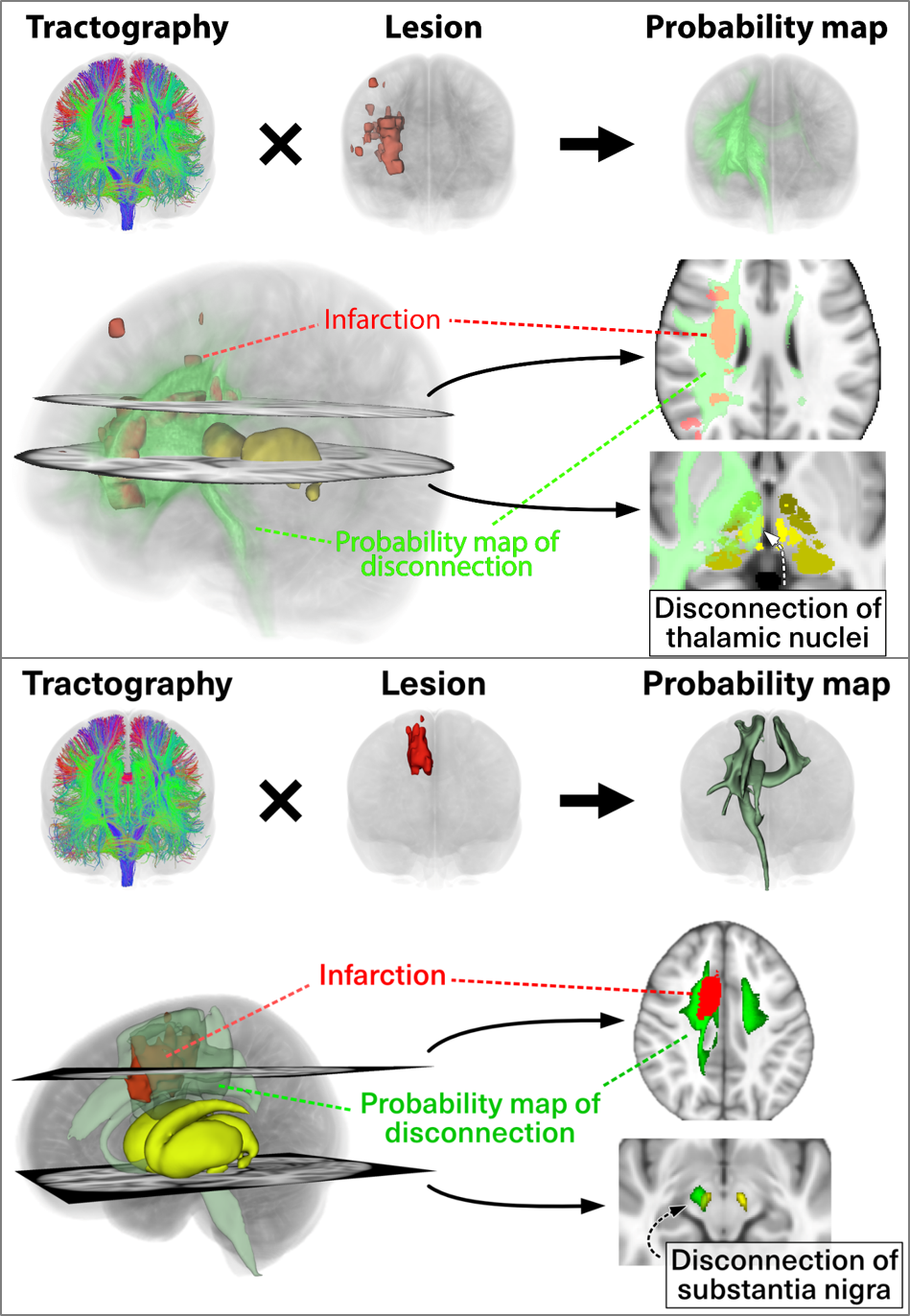

We calculated the percentage of overlap volume between the disconnectivity maps and the regions of interest ROI, which include the four thalamic groups and the substantia nigra, as follows:

$$Overlap \left( \% \right)=\frac{Volume of overlap \left( diconnectivity \subset ROI \right) x probablity of disconnection}{Volume ROI}X 100$$

The overlap histograms for the thalamic nuclei and substantia nigra displayed a left skewed distribution, in which overlap of less than 2.5% categorized the region as “not disconnected. The distribution of the remaining percentage of overlap allowed to dichotomize “mildly disconnected” or “severely disconnected” cases using the median as a cutoff. **Supp-Figure-3** illustrates the distribution of overlap within the medial thalamic group. Similar analyses were conducted for the overlap with the lateral and posterior thalamic groups as well as with the substantia nigra. The anterior thalamic group was too small for accurate disconnection estimation using this methodology and was therefore excluded from further analysis.

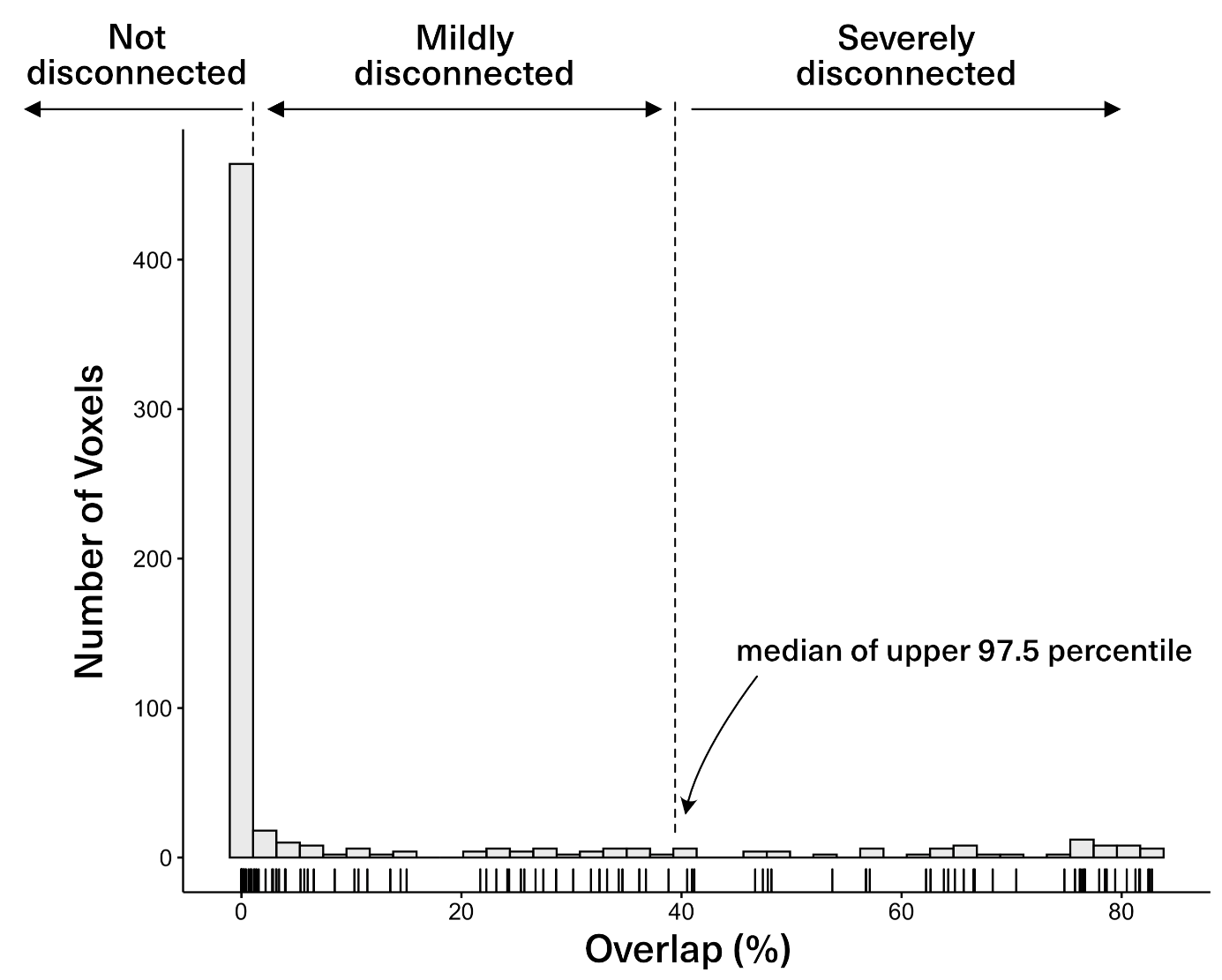

**Supp-figure 3**: Histogram of voxel overlap between the disconnected fibers and the medial thalamic nuclei group among the entire population, and identification of the threshold to consider the nucleus as “not disconnected”, “mildly disconnected” and “severely disconnected”.

**Animal model**

Mice were anesthetized using 4% isoflurane in 20% oxygen, and anesthesia was maintained at 2% isoflurane in 20% oxygen throughout the procedure. After shaving their heads, the mice were positioned in a stereotaxic frame with lidocaine-coated ear bars to minimize movement during surgery. Body temperature was maintained at 37°C using a heated blanket, and eye protection was provided with a gel (Ocry-gel, TVM, Lempdes, France) to prevent drying.

A small incision was made in the scalp to visualize the bregma and lambda, allowing precise positioning of the light source (Schott KL 1600 LED, 680 lumen, green filter, 150 Watts, 5-mm light aperture) at 0 mm antero-posterior and 2.2 mm lateral to the bregma. Rose bengal (Aldrich Chemical Company, Milwaukee, USA) was administered intraperitoneally at a dose of 10 µL/g body weight, with a concentration of 5 mg/mL in saline solution at room temperature. After a 5-minute delay, the light was turned on and directed onto the cranial bone for 10 minutes. Upon completion, the scalp was irrigated and sutured, anesthesia was gradually withdrawn, and the mice were placed in a heated recovery box. Sham procedures were identical, except for the omission of light exposure.

Animals were divided into three groups as detailed in **Supp-Figure-4**

**In vivo MRI acquisition and image analyses in animals**

MRI images were acquired on a 7 Tesla Bruker Biospec system (Ettlingen, Germany) equipped with a gradient system capable of 660 mT/m maximum strength and 110μs rise time. A volume resonator (75.4 mm inner diameter, active length 70 mm) operating in quadrature mode was used for excitation and a 4-element (2 × 2) phased array surface coil (outer dimensions of one coil element: 12 × 16 mm^2^, total outer dimensions: 26 × 21 mm^2^) was used for signal reception.

First, a 2D diffusion sequence was acquired with the following parameters: 82 slices; repetition time, 3000 ms; echo time, 24 ms; slice thickness, 500 µm; matrix, 128 x 80; field of view, 25 mm x 16 mm; b values, 0 and 1000 s/mm^2^; 6 directions; 4 segments; total acquisition time, 5 min 36 s.

A 3D-TrueFISP composite sequence was acquired in 8 separate sequences only differing by their RF phase advance (0°, 45°, 90°, 135°, 180°, 225°, 270° and 315°). Other parameters were the same and as follows: 112 slices; repetition time, 6 ms; echo time, 3 ms; flip angle, 27.5°; slice thickness, 145 µm; matrix, 112 x 112; field of view, 16.2 mm x 16.2 mm; total acquisition time = 12 min 15 s.

To allow R2* quantification, a 3D Multi-Echo T2*-weighted sequence was acquired with the following parameters: 112 slices; repetition time, 64.4 ms; 16 echo times with first echo, 1.61 ms; last echo, 46.61 ms and with a delta-TE of 3 ms; flip angle, 15°; slice thickness, 145 µm; matrix, 112 x 112; field of view, 16.2 mm x 16.2 mm; three averages, total acquisition time = 40 min 23 s.

We extracted R2* related metrics in specific thalamic nuclei and their volumes. For that purpose, we first applied motion correction to the multi-echo T2* images to minimize the influence of movement during image acquisition. Then, the first volume of the multi-echo images was registered to the 3D-TrueFISP using ANTs (12). The anatomical 3D-TrueFISP images were subsequently registered to the template space of a detailed mouse thalamic atlas (13) using Elastix, a robust image registration toolbox (14).

R2* maps were generated from voxel-by-voxel mono-exponential fitting of the motion-corrected and registered multi-echo T2* images and we quantified the 95^th^ percentile of the asymmetry index (AI_95_) in specific nuclei similarly to what we did in patients.

Next, an inverse transformation was applied to project back the thalamic nuclei regions from the atlas space into the native 3D-TrueFISP space. Finally, volumes of the specific nuclei were calculated using the segmented regions obtained from the inverse transformation, and were expressed as asymmetry index (AI) compared to the volumes of contralateral nuclei.

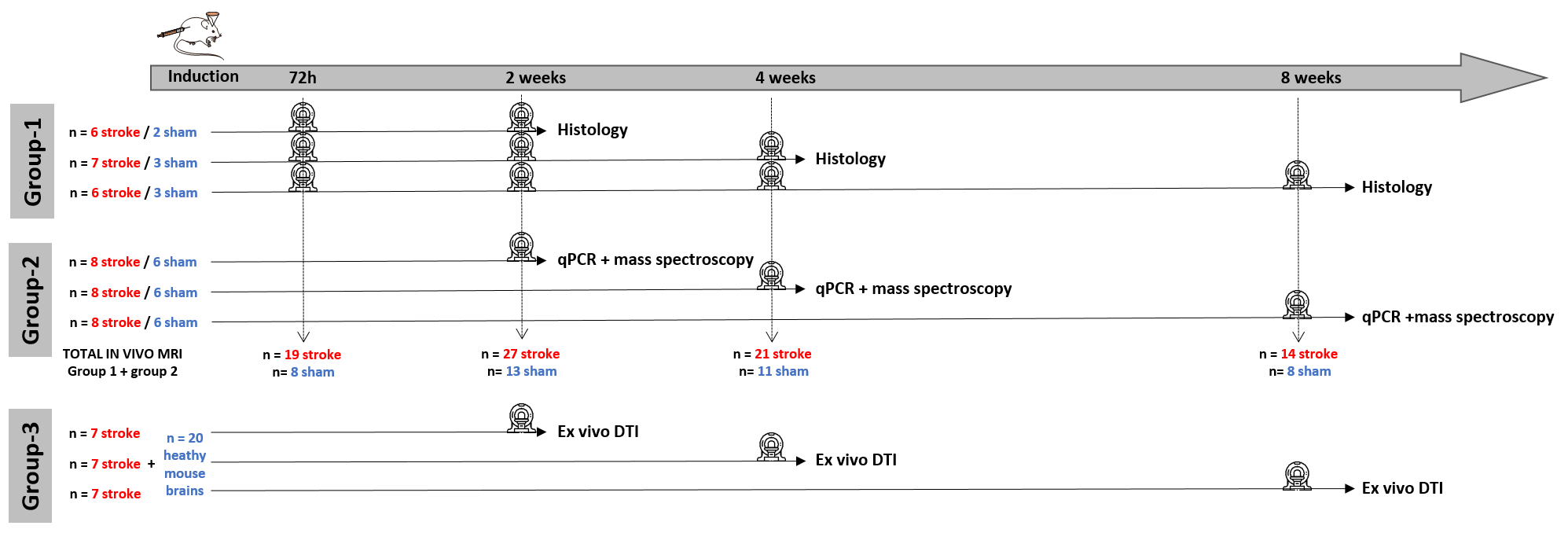

**Supp-Figure 4:** Experimental design indicating the different groups. The pictograms along the timeline indicate magnetic resonance imaging sessions, either *in vivo* for the groups 1 and 2, or *ex vivo* for the group 3.

**Biological analyses**

**Immunohistochemistry**

Mice from the group-1 used for histological analyses were deeply anesthetized with Exagon and lidocaine and perfused transcardially with saline followed by PBS solution containing 4% paraformaldehyde solution, for 8 min. The brains were then removed and transferred into a 4% paraformaldehyde solution, for post-fixation. They were then transferred into a Tris-buffered saline (TBS) containing 30% sucrose and 0.05% sodium azide, to allow cryoprotection, frozen in OCT (Tissue-Tek®, Sakura Finetech, USA) and stored at -80°C until use. A tissue block containing the thalamus was then cut into 30 µm thick coronal sections with a cryostat (Leica CM 1950, Nussloch, Germany).

Immunostaining was performed for microglia (anti-Iba1), astrocytes (anti-GFAP), neurons (anti-NeuN) and ferritin (anti-ferritin) using single and double (Iba1+ferritin, Iba1 + NeuN) staining. Free-floating sections were rinsed in TBS and incubated overnight in a TBS solution containing 0.25% Triton X-100, 1% normal donkey serum and diluted primary antiserum. The following antisera were used: goat anti-Iba1 (1:2000; ABCAM, Ab5076), rabbit anti-Iba1 (1:1000; Sobioda, W1W019-19741), rabbit anti-GFAP (1:2000; DAKO, Z0334), chicken anti-NeuN (1:1500; MERCK, ABN91) and rabbit anti-ferritin light chain (1:800; ABCAM, Ab69090). Afterwards, sections were washed in TBS and incubated using the secondary antibodies (donkey IgGs): anti-rabbit Alexa Fluor 488 (1/1000; Jackson, 711-545-152), anti-goat TRITC (1/1000; Jackson, 705-025-147), anti-rabbit TRITC (1/1000; Jackson, 711-025-152) or anti-chicken TRITC (1/1000; Jackson, 703-025-155) in a TBS solution containing 0.25% Triton X100 (2 h at room temperature). A mounting medium containing DAPI was used for coverslipping (Vectashield H-1200).

Sections were imaged for quantitative analyses on NanoZoomer wide field microscope (Hamamatsu nanozoomer 2.0HT) equipped with a 40x objective and a CCD camera. This system was driven by Mercator software and used to scan slides (mosaic images) for direct comparison with MRI. Identical laser intensity and exposure settings were applied to all tissue sections. At least 2 tissue sections containing the thalamus were randomly selected and quantitative results were averaged. Images were analyzed using ImageJ software. Regions of interest (ROI) outlining thalamic VPL and VPM nuclei were manually drawn using atlas to guide the delineation. Staining areas were quantified automatically as percent of immunoreactivity after subtracting background and thresholding identically all the images for conversion to binary masks.

**Laser capture microdissection**

The mice from group-2 were deeply anesthetized with Exagon and lidocaine and perfused transcardially with PBS for 1 min 30 s. Their brains were quickly extracted, flash-frozen in a vial of isopentane that was immersed in dry ice and subsequently stored at -80 °C until sectioning. Brains were cut using 50 µm thick coronal sections on a freezing microtome (CM3050 S Leica) at -22 °C to prevent RNA degradation. Tissue sections were mounted on PENmembrane 1 mm glass slides (P.A.L.M. Microlaser Technologies AG, Bernried, Germany) that had been pretreated to inactivate RNase. Frozen sections were fixed in a series of precooled ethanol baths (40 s in 95%, 75% and 30 s in 50%) and stained with cresyl violet 1% for 20 s. Subsequently, sections were dehydrated in a series of precooled ethanol baths (30 s in 50%, 75% and 40 s in 95% and in anhydrous 100% twice). Immediately after dehydration laser capture microdissection was performed with a P.A.L.M MicroBeam microdissection system version 4.0-1206 equipped with a P.A.L.M. RoboSoftware (P.A.L.M. Microlaser Technologies AG, Bernried, Germany). Laser power and duration were adjusted to optimize capture efficiency. Microdissection was performed at 5x magnification. The whole thalamus was captured to maximize the amount of material and because isolation of specific nuclei was very difficult to conduct in such conditions. The material was split in 2 adhesive caps (P.AL.M Microlaser Technologies AG, Bernried, Germany) with a 1 slice for qPCR every 3 slices for mass spectrometry.

**Quantitative real-time PCR (qPCR)**

For RNA analysis, we added 150 µl of extraction buffer provided in a ReliaPrep^TM^ RNA Cell Miniprep System (Promega) to the caps and stored at -80 °C until RNA isolation. Total RNA was extracted from microdissected tissues by using the ReliaPrep^TM^ RNA Cell Miniprep System (Promega, Madison, USA) according to the manufacturer’s protocol. The concentration of RNA was determined with Nanodrop 1000. RNA qualities were performed using Agilent RNA 6000 Pico Kit on 2100 Bioanalyzer (Agilent Technologies, Santa Clara, CA).

RNA was processed and analyzed according to an adaptation of published methods (15). Briefly, cDNA was synthesized from 90 ng of total RNA by by using Maxima Reverse Transcriptase (Thermo Scientific). qPCR was performed with a LightCycler® 480 Real-Time PCR System (Roche, Meylan, France). qPCR reactions were done in duplicate for each sample by using transcript-specific primers, cDNA (2.6 ng) and LightCycler 480 SYBR Green I Master (Roche) in a final volume of 10 µl. The qPCR data were exported and analyzed in an informatics tool (Gene Expression Analysis Software Environment) developed at our institute. For the determination of the reference genes, the RefFinder method was used (16). Relative expression analysis was normalized against two reference genes. In particular, peptidylprolyl isomerase A (Ppia) and non-POU-domain-containing, octamer binding protein (Nono) were used as reference genes here. To estimate the PCR amplification efficiency, standard curves were determined, and all the primers chosen were 100% efficient. The relative level of expression was calculated with the comparative (2^_ΔΔCT^) method (17). The following primer sequences were used

| Table X. Primer Sequences. | | |  |  |
| --- | --- | --- | --- | --- |
| **Gene** | **GenBank ID** | **Forward Sequence (5′-3′)** | | **Reverse Sequence (5′-3′)** |
| Itgb2 (CD18) | NM_008404 | TCGGTTTCTTTCCGCCATTA | | AAGATTGTGCAGGTCGGAAGAC |
| Itgam (CD11b) | NM_001082960 | CTCATCACTGCTGGCCTATACAA | | GCAGCTTCATTCATCATGTCCTT |
| Hsbp1 | U03560 | ACGAAGAAAGGCAGGACGAA | | ACCTGGAGGGAGCGTGTATTT |
| Nono | NM_023144 | CTGTCTGGTGCATTCCTGAACTAT | | AGCTCTGAGTTCATTTTCCCATG |
| Ppia | NM_008907 | CAAATGCTGGACCAAACACAA | | GCCATCCAGCCATTCAGTCT |
| Ftl1 | NM_010240 | GGACCCTCATCTCTGTGACTTCC | | GCCCATCTTCTTGATGAGTTTCA |

**Mass spectrometry**

For chemically iron concentration, we used inductively coupled mass spectrometry.

All samples were handled with care in order to avoid environmental contamination during their manipulation.

Brain tissue samples, stored at -80°C, were desiccated at 120°C for 15 hours in an oven. Then, dried samples were weighed and mineralized by nitric acid solution in Teflon PFA-lined digestion vessels. Acid digestion was carried out at 180°C using ultrapure concentrated HNO3 (69%) (Fisher Chemical Optima Grade) in microwave oven device (Mars 6, CEM®).

Iron was measured by Inductively Coupled Plasma Mass Spectrometry (ICP-MS), on an ICAP-TQ from Thermo Scientific® equipped with collision cell technology. The source of plasma was argon (Messer®) with high degree of purity (>99.999%). The collision/reaction cell used was pressurized with helium (Messer®). All rinse, diluent, and standards were prepared with ≥18 MΩ cm ultrapure water using a Millipore® MilliQ Advantage A10 water station.

Digested samples were diluted 1:20 by addition of a diluent solution consisting of 0.5% (v/v) HNO_3_ (Optima Grade Fischer Chemical®). The internal standard used was rhodium (Fisher Scientific®). Calibration ranges preparation was carried out using a multi-element calibrator solution (SCP Science® Plasma Cal). Calibration and verification of instrument performance were realized using multi-element solutions (Thermo®). Quality control was Clincheck controls for trace elements (Recipe). The accuracy of the ICP-MS assay method was verified through analysis of certified reference materials (NCS ZC71001) and participation in External Quality Assessment Schemes (EQAS) organized by OELM (Occupational and Environmental Laboratory Medicine).

**Ex vivo Image acquisition and analyses in animals**

Two groups of ex vivo mouse brains were used:

- 20 healthy mice to build a bilateral template of a thalamo-cortical tract connecting the infarct region to the specific altered thalamic nuclei;

- 21 stroke mice to explore the white matter microarchitecture changes along this thalamo-cortical tract at 2-, 4-, and 8-weeks post-stroke (n=7 per group).

**Healthy animal preparation and** **diffusion MRI acquisition**

The healthy mice were deeply anesthetized at 8 weeks and perfused transcardially with saline followed by PBS solution containing 4% paraformaldehyde solution. Brains were extracted, postfixed in 4% paraformaldehyde overnight and placed on a custom 3D-printed apparatus to prevent movement during the MR acquisition. Each brain in a 3D-printed apparatus was then put into a transparent falcon (1.5mL) and immersed for two days in a proton-free solution of Fomblin YVAC L 14/6 (Kurt J. Lesker) before imaging.

Images were acquired using the 7T Bruker Biospec 70/20 system (Ettlingen, Germany) scanner with a standard Bruker cross coil setup with a quadrature surface coil for mice. Diffusion imaging was acquired using a 3D diffusion-weighted Spin-Echo EPI sequence with the following parameters: 82 slices; repetition time, 1000 msec; echo time, 30 msec; slice thickness, 121µm; matrix, 170 x 110; field of view, 18 mm x 13 mm resulting in a native image resolution of 105 x 118 x 121 µm^3^; b-values, 0 and 1500 s/mm^2^; 60 unique diffusion directions and five non-diffusion direction (b0) measurements; total acquisition time, 9 hours.

**Diffusion MRI data processing**

Diffusion-weighted data pre-processing and analysis were performed with a combination of tools, namely FSL (18), MRtrix3 (19), ANTs (12), and DIPY-based SCILPY (20) [(github.com/scilus/scilpy](http://github.com/scilus/scilpy)). First, raw MRI files were converted to the standard NifTi format and compressed. Next, preprocessing steps were carried out: b0 images were extracted, averaged, and used to create a mask of the whole brain; homogeneity and eddy current corrections were carried out. The averaged b0 volume was aligned to the Allen Mouse Brain Atlas (AMBA) (21) at the 70 µm isotropic resolution using ANTs. Both linear and nonlinear matrices were then used to align the native DWI data in the AMBA space. The fractional anisotropy (FA) maps were then computed.

Tractography was carried out using a constrained spherical deconvolution local tracking algorithm (22) with the following parameter (theta = 20°, step size = 0.05 mm, min/max length 2/20 mm, number of seeds/voxels = 500 in the seeding mask). The seeding mask corresponded to the concatenated white matter ROIs of the AMBA, while the tracking mask was the whole brain mask computed from the averaged b0 volume.

**Extracting a thalamo-cortical tract and building a bilateral template**

Each whole brain tractogram was filtered to extract the streamlines connecting the infarct region to the specific thalamic nuclei in the right hemisphere. First, the cortical infarct regions induced *via* photothrombotic ischemia (n=12 out of the 21 stroke mice) were manually delineated and aligned with the AMBA space, resulting in a single averaged fronto-parietal infarct region. Second, the right thalamic nuclei in AMBA, specifically the ventral posterolateral (VPL) and ventral posteromedial (VPM) nuclei, regions affected by the induced stroke, were selected as the second filtering ROI. We concatenated the 20 thalamo-cortical tracts to create the right hemisphere's final template (Figure 6A, top). Given the symmetrical structure of the AMBA, this right-hemisphere template was mirrored across the interhemispheric plane to generate a corresponding template for the left hemisphere.

**Fractional anisotropy measurements**

We used a tract profile approach to estimate the FA values along the thalamo-cortical template in the stroke mice at 2-, 4-, and 8-weeks post-stroke (7 mice per time point). The label image of the thalamo-cortical template represented the coverage of the segmented tract into regions labeled from 1 to 20, starting from the thalamic nuclei (Figure 6A, bottom). We then calculated the mean (± standard deviation) FA values within each labeled region for each stroke mouse at each post-stroke duration (Figure 6B).

**Supplemental results**

**Supplemental results on substantia nigra disconnection in patients**

**Supp-Table 1** shows the characteristics of the patients included in the substantia nigra analysis.

| Variables | Participants included in the substantia nigra analysis (n=177) |
| --- | --- |
| Demographics |  |
| Age (y)* | 65 (56, 77) |
| Sex |  |
| M | 120 (68) |
| F | 57 (32) |
| Hypertension | 112 (63) |
| Diabetes mellitus | 28 (16) |
| Active smoking | 83 (47) |
| Body mass index (Kg/m²)* | 26.7 (24.2, 29.3) |
| Baseline visit |  |
| NIHSS score* | 3.0 (2.0, 6.0) |
| Time from onset to baseline MRI (h)* | 47 (33, 60) |
| Recanalization procedure (thrombolysis and/or thrombectomy) | 85 (48) |
| Infarct volume (mL)* | 10 (2, 29) |
| Follow-up visit |  |
| Time from onset to follow-up (y)* | 1.0 (0.98, 1.01) |
| NIHSS score* | 0.5 (0.0, 2.0) |
| Modified Rankin scale (mRS)* | 1.0 (0.0, 2.0) |

Except where indicated, data are numbers of patients with percentages in parentheses. * Data are median, with IQRs in parentheses..

**Supp-Table 2:** characteristics of the population according to the disconnectivity status for the lateral, medial and posterior thalamic nuclei groups and for the substantia nigra.

|  | Lateral group of thalamic nuclei | | | Medial group of thalamic nuclei | | | Posterior group of thalamic nuclei | | | Substantia nigra | | |
| --- | --- | --- | --- | --- | --- | --- | --- | --- | --- | --- | --- | --- |
|  | Not disconnected | Mildly disconnected | Severely disconnected | Not disconnected | Mildly disconnected | Severely disconnected | Not disconnected | Mildly disconnected | Severely disconnected | Not disconnected | Mildly disconnected | Severely disconnected |
|  | n=50 | n=53 | n=53 | n=79 | n=38 | n=39 | n=60 | n=48 | n=48 | n=54 | n=61 | n=62 |
| Age (y)* | 67 (57 – 77) | 63 (54 – 71) | 66 (60 – 77) | 66 (57 – 77) | 62 (51 – 76) | 66 (60 – 75) | 66 (56 – 78) | 63 (57 – 74) | 65 (57 – 74) | 68 (56 – 77) | 63 (55 – 73) | 66 (58 – 76) |
| M / F | 38 (76) / 12 (24) | 40 (75) / 13 (25) | 30 (57) / 23 (43) | 60 (76) / 19 (24) | 25 (66) / 13 (34) | 23 (59) / 16 (41) | 45 (75) / 15 (25) | 35 (73) / 13 (27) | 28 (58) / 20 (42) | 42 (78) / 12 (22) | 41 (67) / 20 (33) | 37 (60) / 25 (40) |
| Hypertension | 26 (52) | 36 (68) | 39 (74) | 51 (65) | 23 (61) | 27 (69) | 32 (53) | 33 (69) | 36 (75) | 30 (56) | 41 (67) | 41 (66) |
| Diabetes mellitus | 5 (10) | 11 (21) | 10 (19) | 12 (15) | 9 (24) | 5 (13) | 8 (13) | 9 (19) | 9 (19) | 5 (9.3) | 12 (20) | 11 (18) |
| Active smoking | 21 (42) | 28 (53) | 23 (43) | 35 (44) | 21 (55) | 16 (41) | 28 (47) | 22 (46) | 22 (46) | 25 (46) | 30 (49) | 28 (45) |
| Body mass index (Kg/m²)* | 26.8 (24.3 – 29.6) | 25.9 (24.0 – 28.3) | 27.7 (24.1 – 30.1) | 26.5 (24.2 – 29.0) | 26.1 (23.7 – 28.4) | 27.7 (25.4 – 30.1) | 25.6 (23.8 – 28.4) | 26.5 (23.8 – 28.4) | 28.3 (26.2 – 30.4) | 26.8 (24.4 – 29.7) | 25.7 (23.9 – 27.8) | 27.7 (24.2 – 30.2) |
| Baseline NIHSS score* | 2 (1 – 3) | 3 (2 – 4) | 7 (4 – 13) | 2 (1 – 3) | 4 (3 – 7) | 6 (3 – 16.5) | 2 (1 – 3) | 3 (2 – 5) | 7 (4 – 13) | 2 (1 – 2) | 3 (2 – 5) | 6 (3 – 13) |
| Baseline Infarct volume (mL)* | 7.90 (1.50 – 16.49) | 3.90 (0.89 – 15.85) | 19.49 (8.31 – 65.92) | 5.89 (1.23 – 13.74) | 4.93 (1.75 – 21.43) | 27.05 (15.32 – 80.58) | 5.60 (1.31 – 13.02) | 9.03 (1.70 – 21.23) | 22.96 (5.85 – 70.96) | 7.90 (1.50 – 17.01) | 2.40 (0.90 – 11.68) | 20.38 (8.75 – 52.34) |
| NIHSS score at follow-up* | 0 (0 – 1) | 0 (0 – 1) | 2 (1 – 6) | 0 (0 – 1) | 1 (0 – 3) | 1 (0.5 – 6) | 0 (0 – 1) | 1 (0 – 1) | 2 (0 – 6) | 0 (0 – 1) | 1 (0 – 2) | 1 (0 – 5) |
| mRS at follow-up* | 0 (0 – 1) | 1 (0 – 1.25) | 2 (1 – 3) | 0 (0 – 1) | 2 (1 – 2) | 1.5 (1 – 3) | 0.5 (0 – 1) | 1 (0 – 2) | 2 (1 – 3) | 0 (0 – 1) | 1 (0.5 – 2) | 2 (0.25 – 3) |

Except where indicated, data are numbers of patients with percentages in parentheses. * Data are median, with IQRs in parentheses.

Participants with severely disconnected substantia nigra showed a significant increase of AI_95_ from baseline to 1 year while there was no modification of AI_95_ if this structure was not or mildly disconnected (**Supp-Figure-5**).

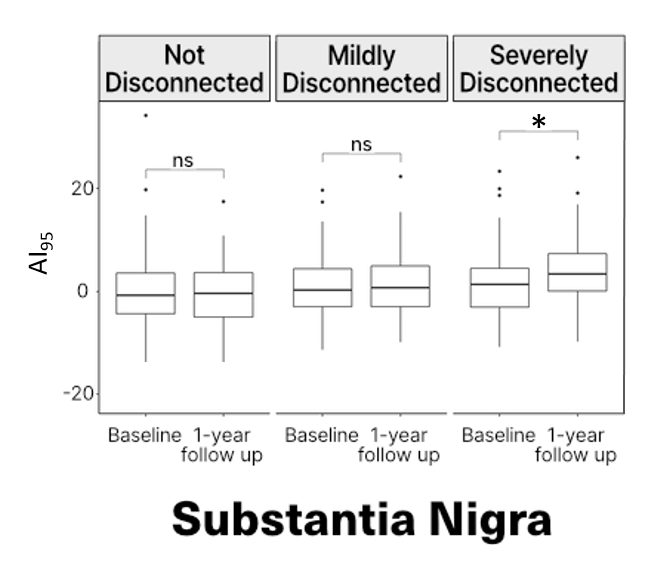

**Supp-Figure 5** shows the relationship between the substantia nigra disconnectivity status and the secondary R2* increase at 1-year follow-up.

The linear regression confirmed that the disconnectivity status of susbtantia nigra at baseline was a significant and independent predictor of AI_95_ at follow-up (**Supp-Table 3**).

**Supp-Table 3:** Multivariable linear regression models for prediction of AI_95_ at 1 year in substantia nigra

| Variable | Β value (95%CI | *P* Value |
| --- | --- | --- |
| Disconnection of the SUBSTANTIA NIGRA | | |
| Age (+1 y) | -0.03 (-0.18, 0.11) | 0.70 |
| Sex (F) | -0.15 (-0.40, 0.17) | 0.40 |
| AI_95_ at baseline (+ 1) | **0.15 (0.01, 0.30)** | **0.042*** |
| Infarct volume (+ 1 mL) | 0.12 (-0.04, 0.27) | 0.13 |
| Disconnection |  |  |
| Mildly disconnected | 0.26 (-0.10, 0.62) | 0.20 |
| Severely disconnected | **0.55 (0.18, 0.92)** | **0.004**** |

**Supplemental results on the photothrombosis stroke model in mice**

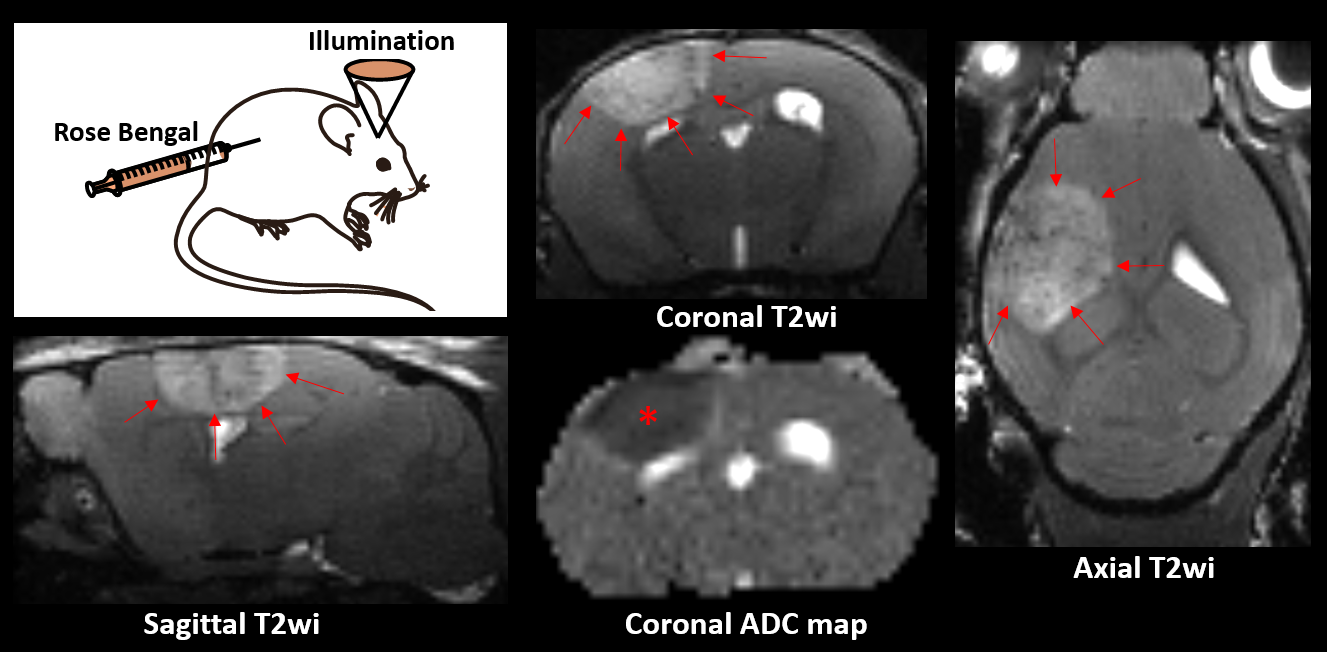
The photothrombosis model induced a reproducible lesion within the sensory motor cortex **(Supp-Figure-6)**

**Supp-Figure 6** shows a schematic representation of the photothrombosis model that consists in transcranial illumination of the brain 5 minutes after intraperitoneal injection of the Rose Bengal photosensitive dye. After a delay of 72h, we verified with MRI that we consistently induced a focal cortical stroke (delineated by arrows on T2 weighted images) that showed low ADC values (*) in line with the cytotoxic edema encountered at the acute stage on an infarct.

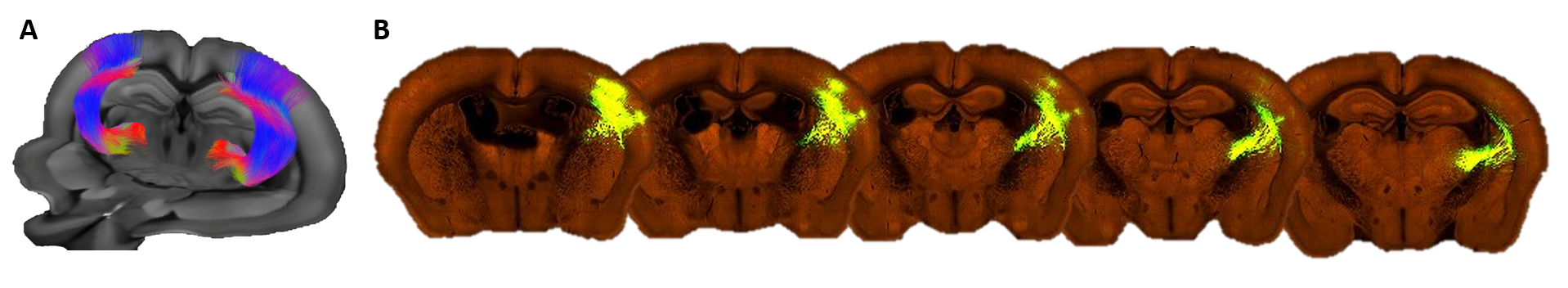
**Supp-Figure 7: (A)** shows the 3D projection of fibers connecting the cortical site to ipilateral VPL and VPM from diffusion tensor MRI. **(B)** shows the close correspondence with serial two-photon tomography after 0.18 mm^3^ EGFP tracer injected in VPL and VPM of Slc17a6-IRES-Cre mice from mouse connectivity data from the Allen Brain Atlas portal (23).

23.	<https://connectivity.brain-map.org/>.
